# Mechanisms underlying serotonergic excitation of callosal projection neurons

**DOI:** 10.1101/197624

**Authors:** Emily K. Stephens, Arielle L. Baker, Allan T. Gulledge

## Abstract

Serotonin (5-HT) selectively excites subpopulations of pyramidal neurons in the neocortex via activation of 5-HT_2A_ (2A) receptors coupled to G_q_ subtype G-protein alpha subunits. Gq-mediated excitatory responses have been attributed primarily to suppression of potassium conductances, including those mediated by K_V_7 potassium channels (i.e., the M-current), or activation of nonspecific cation conductances that underly calcium-dependent afterdepolarizations (ADPs). However, 2A-dependent excitation of cortical neurons has not been extensively studied, and no consensus exists regarding the underlying ionic effector(s) involved. We tested potential mechanisms of serotonergic excitation in commissural/callosal projection neurons (COM neurons) in layer 5 of the mouse medial prefrontal cortex, a subpopulation of cortical pyramidal neurons that exhibit 2A-dependent excitation in response to 5-HT. In baseline conditions, 5-HT enhanced the rate of action potential generation in COM neurons experiencing suprathreshold somatic current injection. This serotonergic excitation was occluded by activation of muscarinic acetylcholine (ACh) receptors, confirming that 5-HT acts via the same G_q_-signaling cascades engaged by ACh. Like ACh, 5-HT promoted the generation of calcium-dependent ADPs following spike trains. However, calcium was not necessary for serotonergic excitation, as responses to 5-HT were enhanced (by >100%), rather than reduced, by chelation of intracellular calcium with 10 mM BAPTA. This suggests intracellular calcium negatively regulates additional ionic conductances contributing to 2A excitation. Removal of extracellular calcium had no effect when intracellular calcium signaling was intact, but suppressed 5-HT response amplitudes, by about 50% (i.e., back to normal baseline values) when BAPTA was included in patch pipettes. This suggests that 2A excitation involves activation of a nonspecific cation conductance that is both calcium-sensitive and calcium-permeable. M-current suppression was found to be a third ionic effector, as blockade of K_V_7 channels with XE991 (10 μM) reduced serotonergic excitation by ∼50% in control conditions, and by ∼30% with intracellular BAPTA present. These findings demonstrate a role for at least three distinct ionic effectors, including K_V_7 channels, a calcium-sensitive and calcium-permeable nonspecific cation conductance, and the calcium-dependent ADP conductance, in mediating serotonergic excitation of COM neurons.

## Introduction

Glutamatergic pyramidal neurons represent a key cellular information processing unit within the cortex. Their activity is regulated by modulatory neurotransmitters such as acetylcholine (ACh), dopamine, norepinephrine, and serotonin (5-HT) that act primarily through G-protein coupled receptors. Depending on receptor-specific coupling to G-protein alpha subunits, modulatory neurotransmitters can enhance or suppress synaptic integration and action potential generation in target neurons. For example, in neocortical pyramidal neurons, serotonergic activation of 5-HT_2A_ (2A) receptors coupled to G_q_ subtype G-protein alpha subunits enhances excitability, while activation of G_i/o_-coupled 5-HT_1A_ (1A) receptors suppresses action potential generation (Avesar and Gulledge, 2012; Stephens et al., 2014). However, while stimulation of specific 5-HT receptors leads to predictable changes in action potential output, the ionic mechanisms underlying serotonergic excitation are less well understood.

While a consensus exists that 1A-dependent inhibition is mediated by enhanced potassium conductance (Andrade and Nicoll, 1987; Colino and Halliwell, 1987; Davies et al., 1987; Tanaka and North, 1993; Spain, 1994; Luscher et al., 1997; Goodfellow et al., 2009), the ionic mechanisms responsible for 2A-dependent serotonergic excitation of neocortical pyramidal neurons have been less well studied (Araneda and Andrade, 1991; Spain, 1994; Villalobos et al., 2005; Villalobos et al., 2011), in part because 2A receptors are not universally expressed in adult pyramidal neurons. As a result, much of our understanding of 2A signaling is inferred from studies of cholinergic excitation, which is mediated by the more ubiquitous G_q_-coupled M1-subtype muscarinic ACh receptor. Classically, G_q_-coupled excitation in pyramidal neurons is associated with suppression of potassium conductances, including those mediating the “M” (muscarine-suppressed) current and those responsible for afterhyperpolarizations (AHPs) in pyramidal neurons following bouts of action potential generation (Krnjevic et al., 1971; Schwindt et al., 1988; McCormick and Williamson, 1989; Araneda and Andrade, 1991; Wang and McCormick, 1993; Villalobos et al., 2005; Villalobos et al., 2011). However, reduction of potassium conductances does not fully account for G_q_-mediated excitation in neocortical pyramidal neurons (Andrade, 1991; Haj-Dahmane and Andrade, 1996), as G_q_-coupled muscarinic ACh and 5-HT receptors also engage inward currents generated by calcium-dependent (Haj-Dahmane and Andrade, 1998; Yan et al., 2009; Lei et al., 2014) and independent (Haj-Dahmane and Andrade, 1996; Shalinsky et al., 2002; Egorov et al., 2003; Magistretti et al., 2004) nonspecific cation channels.

Here, to better understand the mechanisms underlying 2A-mediated excitation in the cerebral cortex, we take advantage of the selective and robust 2A excitation that occurs in a subpopulation of pyramidal neurons that project to the contralateral cerebral hemisphere (Avesar and Gulledge, 2012). Using fluorescent retrograde tracers injected unilaterally in the mouse medial prefrontal cortex (mPFC), we challenged 2A excitation in commissural/callosal (COM) projection neurons with a combination of pharmacological and ionic substitution approaches. Our results demonstrate that serotonin 2A receptors engage multiple ionic effectors, including the M-current, a calcium-sensitive but calcium-permeable nonspecific cation conductance, and the calcium-dependent ADP nonspecific cation conductance, to promote action potential generation in COM neurons.

## Materials and Methods

### Animals

Experiments involved adult (6-to-10-week-old) C57BL/6J male and female mice housed in 12:12 hour light:dark cycle and provided food and water *ad libitum*. All experiments were carried out according to methods approved by the Institutional Animal Care and Use Committee of Dartmouth College. No significant sex-dependent differences in cellular physiology or 5-HT responses were observed (**Table 1**).

**Table 1.** Physiology and serotonin responses in neurons from female and male mice.

| Sex | n | Physiology |  |  | Serotonin responses |  |
| --- | --- | --- | --- | --- | --- | --- |
|  |  | RMP<br>(mV) | R<sub>N</sub><br>(MΩ) | I<sub>h</sub><br>(% sag) | % Increase in<br>firing rate | Response<br>integral (Hz·s) |
| **Female** | 15 | -80 ± 5 | 176 ± 46 | 8 ± 4 | 109 ± 60 | 112 ± 92 |
| **Male** | 33 | -80 ± 5 | 181 ± 69 | 9 ± 5 | 85 ± 41 | 129 ± 111 |
| **p-value** |  | 0.67 | 0.79 | 0.74 | 0.11 | 0.61 |
| **All** | 48 | -81 ± 5 | 171 ± 63 | 9 ± 4 | 92 ± 48 | 124 ± 105 |

Data presented as means ± standard deviations.

### Retrograde labeling

Red or green fluorescent microbeads (Retrobeads, Lumafluor Inc.) were injected unilaterally into the left prelimbic cortex using age-appropriate coordinates (Paxinos and Franklin, 2004) to label COM neurons in the contralateral mPFC. Animals were anesthetized throughout surgeries with vaporized isoflurane (∼2%). Following craniotomy, a microsyringe was lowered into the prefrontal cortex and 300 nL of undiluted Retrobead solution was injected over a 10 minute period. Animals were allowed to recover from surgery for at least 72 hours before use in electrophysiological experiments. The location of dye injection was confirmed *post hoc* in coronal sections of the mPFC.

### Slice preparation

Following isoflurane anesthesia and decapitation, brains were removed and submerged in artificial cerebral spinal fluid (aCSF) containing, in mM: 125 NaCl, 25 NaHCO_3_, 3 KCl, and 1.25 NaH_2_PO_4_, 0.5 CaCl_2_, 5 MgCl_2,_ 25 glucose, and saturated with 95% O_2_ / 5% CO_2_. Coronal brain slices (250 μm thick) of the mPFC were cut using a Leica VT 1200 slicer and stored in a chamber filled with aCSF containing 2 mM CaCl_2_ and 1mM MgCl_2_ at 35°C for ∼45 minutes, then kept at room temperature (∼26 °C) for up to 8 hours prior to use in experiments.

### Electrophysiology

Slices were transferred to a recording chamber continuously perfused with oxygenated aCSF at 35-36 °C and visualized with an Olympus BX51WI microscope. Whole-cell current-clamp recordings of layer 5 pyramidal neurons (L5PNs) were made with patch pipettes (∼5 MΩ) filled with, in mM: 135 K-gluconate, 2 NaCl, 2 MgCl_2_, 10 HEPES, 3 Na_2_ATP, and 0.3 NaGTP (pH 7.2 with KOH). Epifluorescence illumination using 470 nm or 530 nm LEDs was used to target Retrobead-labeled layer 5 COM neurons in the prelimbic cortex for whole-cell recording. Data were acquired with Axograph software (Axograph Scientific) using a BVC-700 amplifier (Dagan Corporation) and an ITC-18 digitizer (HEKA Instruments). Membrane potentials were sampled at 25 kHz, filtered at 10 kHz, and corrected for a junction potential of +12 mV.

5-HT (100 μM) was dissolved in aCSF and loaded into a patch pipette placed ∼50 μm from the targeted soma. Neurons were classified as “COM-excited” or “COM-biphasic” based on their initial response to 5-HT (delivered for 1 s at ∼10 PSI) during periods of action potential generation (∼6 Hz) driven by somatic DC current injection (see Avesar and Gulledge, 2012). To preserve focus on 2A-driven signaling, only COM-excited neurons were used for experiments. Serotonergic excitation was quantified as the average instantaneous spike frequency (ISF) over the first 500 ms following the peak post-5-HT increase in ISF, as measured relative to the mean baseline ISF (ie., as the percent increase above baseline firing rates), and as the integral of the increased ISF over time.

### Pharmacological manipulations

Carbachol (50-100 μM; Sigma Aldrich) was used to activate muscarinic receptors. XE991 (10 μM; Tocris Bioscience or Caymen Chemicals) was used to block K_V_7 channels. 1,2-Bis(2-Aminophenoxy)ethane-N,N,N′, N′-tetraacetic acid (BAPTA, 10 mM; Sigma Aldrich) was included in the patch-pipette solution in some experiments to chelate intracellular calcium. For experiments using nominally calcium-free aCSF, CaCl_2_ was replaced with either equimolar or 5 mM MgCl_2_ (no differences were observed between these two conditions). In some experiments, extracellular KCl was reduced to 0.5 mM, with NaCl being raised to 127.5 mM.

### Statistical analyses

Unless otherwise noted, data are presented as mean ± SEM, and were assessed with either Student’s t-test (two-tailed, paired or unpaired) or one-way ANOVA (two-tailed, repeated measures where appropriate) using Wizard for Mac version 1.9 (Evan Miller). Significance was defined as *p* < 0.05.

## Results

### Cholinergic activation of L5PNs occludes serotonergic excitation

To characterize the ionic mechanisms underlying 2A-receptor-mediated serotonergic excitation in the cerebral cortex, we made electrical recordings from labeled callosal/ commissural (COM) projection neurons in layer 5 of the mouse mPFC. Focal application of 5-HT (1 s at ∼10 PSI) during current-evoked action potential generation (∼6 Hz; **Figure 1A**) produced an increase in ISF of 117 ± 23% (n = 9; p < 0.001, paired Student’s t-test) relative to baseline firing frequencies, with response integrals being 168 ± 34 Hz•s (**Figure 1C**). Since these serotonergic responses were qualitatively similar to cholinergic responses mediated by M1 muscarinic ACh receptors (Araneda and Andrade, 1991; Haj-Dahmane and Andrade, 1999; Gulledge et al., 2009), we tested whether these two G_q_-mediated responses share common intracellular signaling cascades by measuring responses to 5-HT before and after tonically activating muscarinic ACh receptors with carbachol (50 - 100 μM, bath applied). Carbachol depolarized COM neurons, and lowered the current necessary to evoke sustained action potential generation by 67 ± 26 pA (54 ± 16%), and induced spontaneous firing in 3/9 neurons. Further, the presence of carbachol fully occluded serotonergic excitation, as no increase in firing rate was observed in response to focally applied 5-HT (n = 9; **Figure 1A**,**B**). Instead, 5-HT application in the presence of carbachol *reduced* the firing frequency of COM neurons (peak change in ISF = -40 ± 10%; n = 9, p = 0.003, paired Student’s t-test; **Figure 1C**), resulting in negative response integrals (-103 ± 39 Hz•s). These results reveal significant overlap of G_q_-signaling cascades downstream of M1 and 2A receptors in COM neurons.

**Figure 1.**
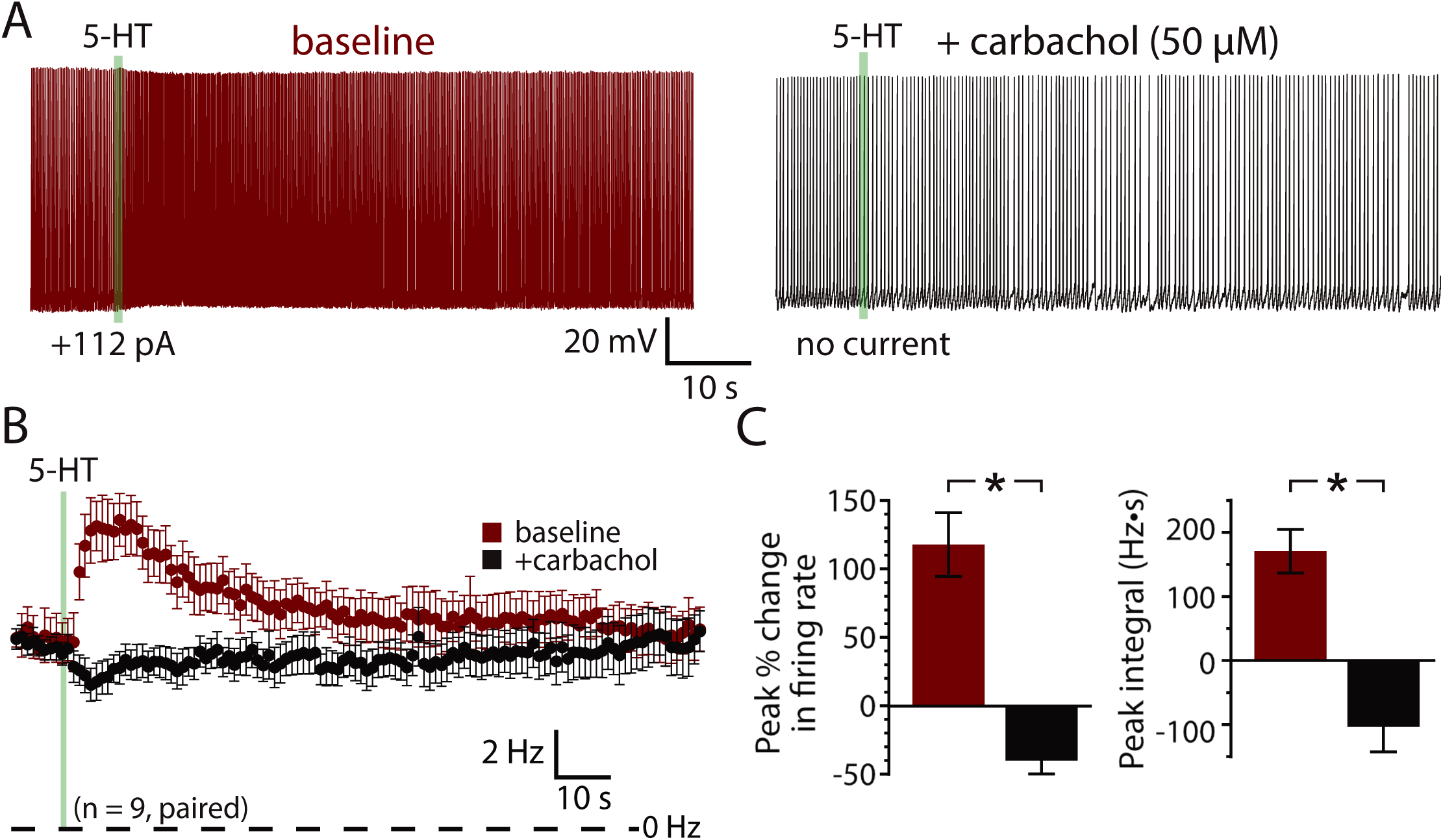
Serotonergic and cholinergic excitation share signaling mechanisms. **(A)** Voltage traces showing responses to focal application of 5-HT (100 μM, 1 s, ∼ 15 psi; green bar) during periods of action potential generation (∼6 Hz) in control conditions (left) and after bath-application of the cholinergic agonist carbachol (right). **(B)** Plots of mean (± SEM) instantaneous spike frequencies in baseline conditions (red) and in the presence of 50 - 100 μM carbachol (black) for 9 neurons. **(C)** Comparisons of the percent change in firing (left) and response integrals (right) for serotonergic responses in control and carbachol conditions.

### 5-HT generates afterdepolarizations (ADPs) in COM neurons

A prominent feature of muscarinic excitation of pyramidal neurons is the generation of afterdepolarizations (ADPs) that occur after pairing M1 receptor activation with brief suprathreshold current injections (Andrade, 1991; Haj-Dahmane and Andrade, 1998; Gulledge et al., 2009; Dasari et al., 2013). Similarly, 5-HT acting via 2A receptors can initiate ADPs in neocortical pyramidal neurons (Araneda and Andrade, 1991; Spain, 1994; Zhang and Arsenault, 2005). We confirmed that 5-HT promotes ADPs in COM neurons by evoking trains of ten action potentials using brief (2 ms) high-amplitude (5 nA) current steps presented at 25 Hz (**Figure 2A**). On some trials, 5-HT was focally applied 1 s before current pulses, and peak depolarization relative to the resting membrane potential (RMP) was measured in the first 1 s following the spike train. Pairing 5-HT with current pulses generated small ADPs with mean amplitudes of 1.6 ± 0.2 mV (n = 16; p < 0.001, paired Student’s t-test; **Figure 2B**). Serotonergic ADPs were absent when the calcium chelating agent BAPTA (10 mM) was included in the patch-pipette solution (n = 6; **Figure 2B**), confirming that they are gated by intracellular calcium signaling (Spain, 1994; Haj-Dahmane and Andrade, 1998; Zhang and Arsenault, 2005; Dasari et al., 2013). These results suggest that the calcium-dependent and G_q_-triggered ADP may represent a common ionic mechanism contributing to serotonergic and cholinergic excitation in COM neurons.

**Figure 2.**
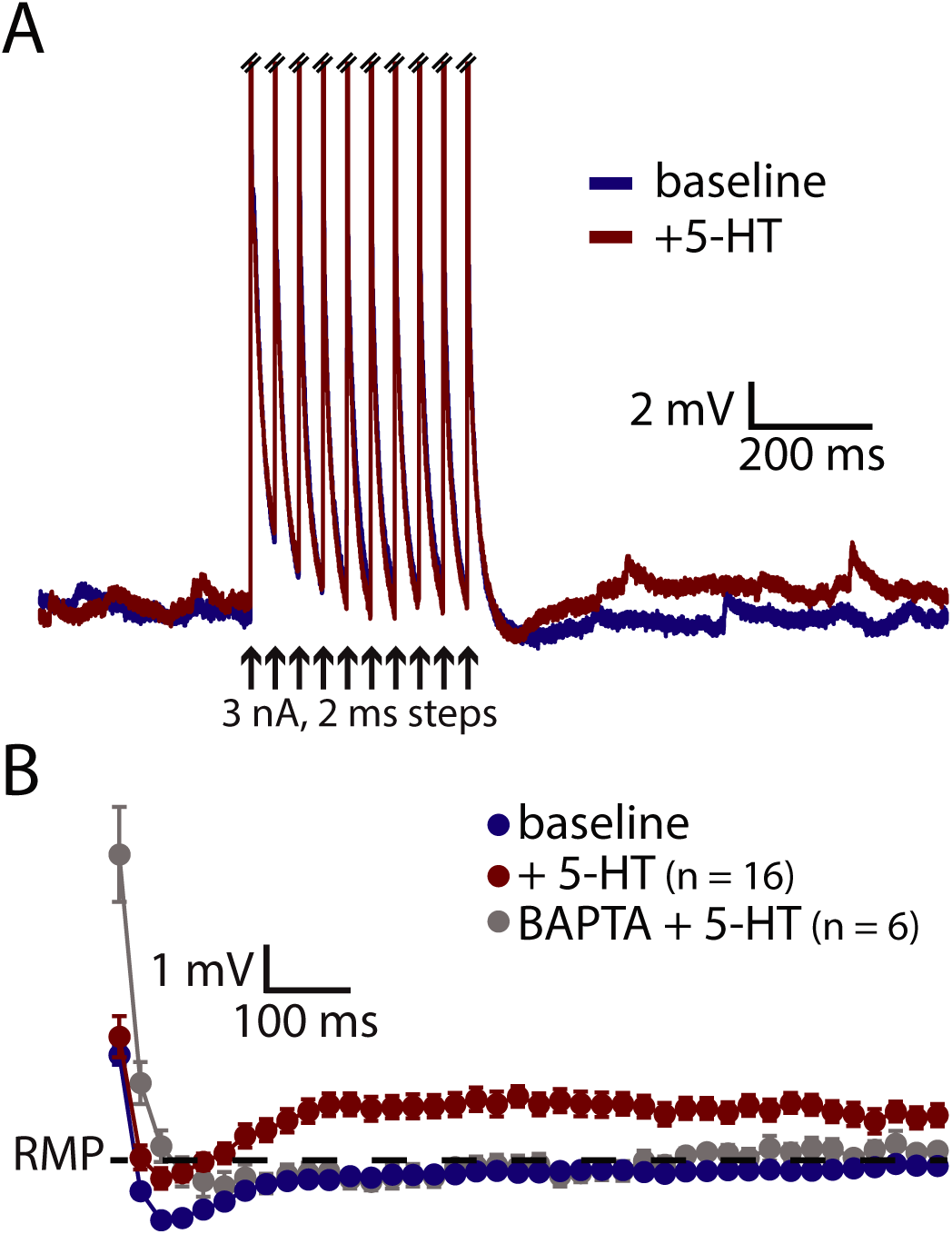
5-HT induces a calcium-dependent afterdepolarization (ADP) in COM neurons. **(A)** Voltage traces from a COM neuron experiencing a train of ten high-amplitude (5 nA, 2 ms, 25 Hz) current injections in baseline conditions (blue) and following focal application of 5-HT (red). Action potentials are truncated. Black arrows indicate current step applications. **(B)** Average voltage responses following spike trains paired with 5-HT, resampled to 50 Hz and plotted as mean ± SEM, for 16 neurons in baseline conditions (blue) and after 5-HT (red). Grey symbols show responses following 5-HT application in a different group of COM neurons filled with 10 mM BAPTA (n = 6).

### Role of calcium in serotonergic excitation of COM neurons

If the ADP current is the major contributor to serotonergic excitation in COM neurons, chelation of intracellular calcium with BAPTA should reduce or eliminate excitatory responses to 5-HT. Instead, addition of BAPTA (10 mM) to the pipette solution (n = 13) enhanced serotonergic excitation, with mean 5-HT response amplitudes being more than twice as large as in COM neurons patched with normal intracellular solution (**Figure 3**). In control neurons (n = 10), peak increases in ISF were 86 ± 11% above baseline values, while in BAPTA-filled neurons peak ISF increases were 226 ± 23% above baseline values (p < 0.001 vs control; Student’s t-test). While in these groups response integrals in BAPTA-filled COM neurons (249 ± 56 Hz•s) were not significantly larger than those in control neurons (147 ± 29 Hz•s; p = 0.13; **Figure 3C**), in three additional independent experimental groups (see below) we observed significant enhancement of both amplitudes and integrals of 5-HT responses in BAPTA-filled neurons relative to controls. These findings demonstrate that serotonergic excitation, as a whole, is not calcium-dependent, and suggest that the calcium-dependent ADP may not represent the major ionic effector mediating 5-HT responses in COM neurons. Instead, the enhancement of responses in the presence of intracellular BAPTA suggests that intracellular calcium acts as a negative regulator for additional ionic effectors contributing to 2A-mediated excitation.

**Figure 3.**
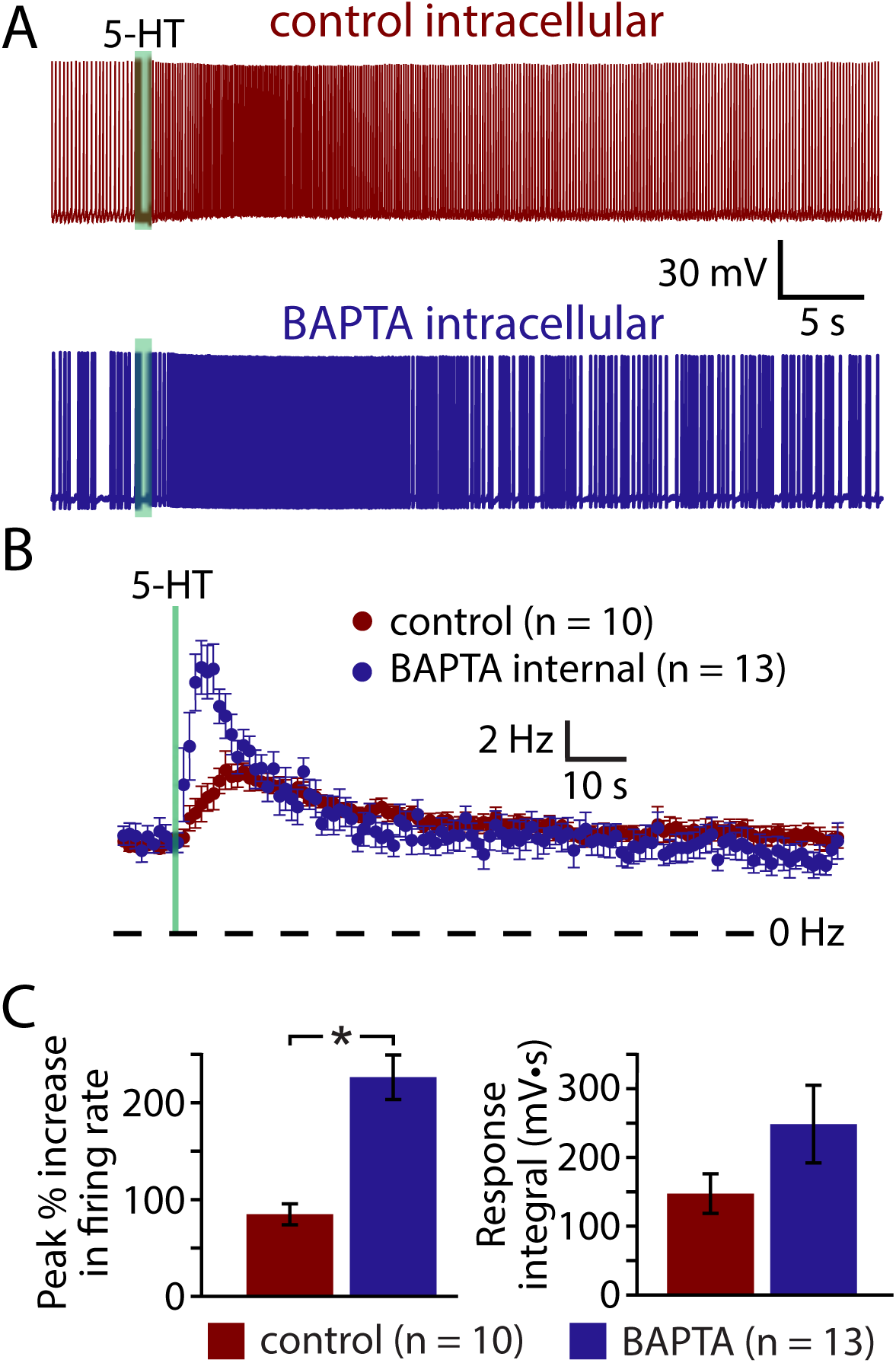
Serotonergic excitation of COM neurons is not calcium-dependent. **(A)** Serotonergic responses in COM neurons recorded with control intracellular solution (red) or with 10 mM BAPTA included (blue). **(B)** Plots of mean (± SEM) instantaneous spike frequencies in control (red; n = 10) and BAPTA-containing (blue; n = 13) COM neurons. **(C)** Comparisons of the peak increase in instantaneous spike frequencies (left) and response integrals (right) for COM neurons recorded in control (red) or BAPTA (blue) conditions.

If activity- and/or 5-HT-driven calcium entry negatively regulates serotonergic excitation of COM neurons, removal of extracellular calcium should enhance 5-HT responses in a manner similar to inclusion of intracellular BAPTA. To test this possibility, we measured responses to 5-HT in COM neurons in baseline conditions and after replacing extracellular calcium with magnesium (see Methods; **Figure 4**). Successful elimination of extracellular calcium was verified in a subset of neurons (n = 4) by measuring electrically evoked EPSPs before, during, and after calcium replacement (**Figure 4A**). While removal of extracellular calcium eliminated synaptic responses, it did not affect the magnitude or integrals of serotonergic responses (n = 9; **Figure 4B**,**C**,**D**). Peak increases in ISFs were 78 ± 11% and 75 ± 12% above initial firing frequencies in baseline and calcium-free conditions, respectively (p = 0.83, paired Student’s t-test), while response integrals for the two conditions were 141 ± 29 Hz•s and 164 ± 44 Hz•s, respectively (p = 0.58). Thus, with normal intracellular calcium signaling intact, removal of extracellular calcium did not affect 5-HT response amplitudes or integrals.

**Figure 4.**
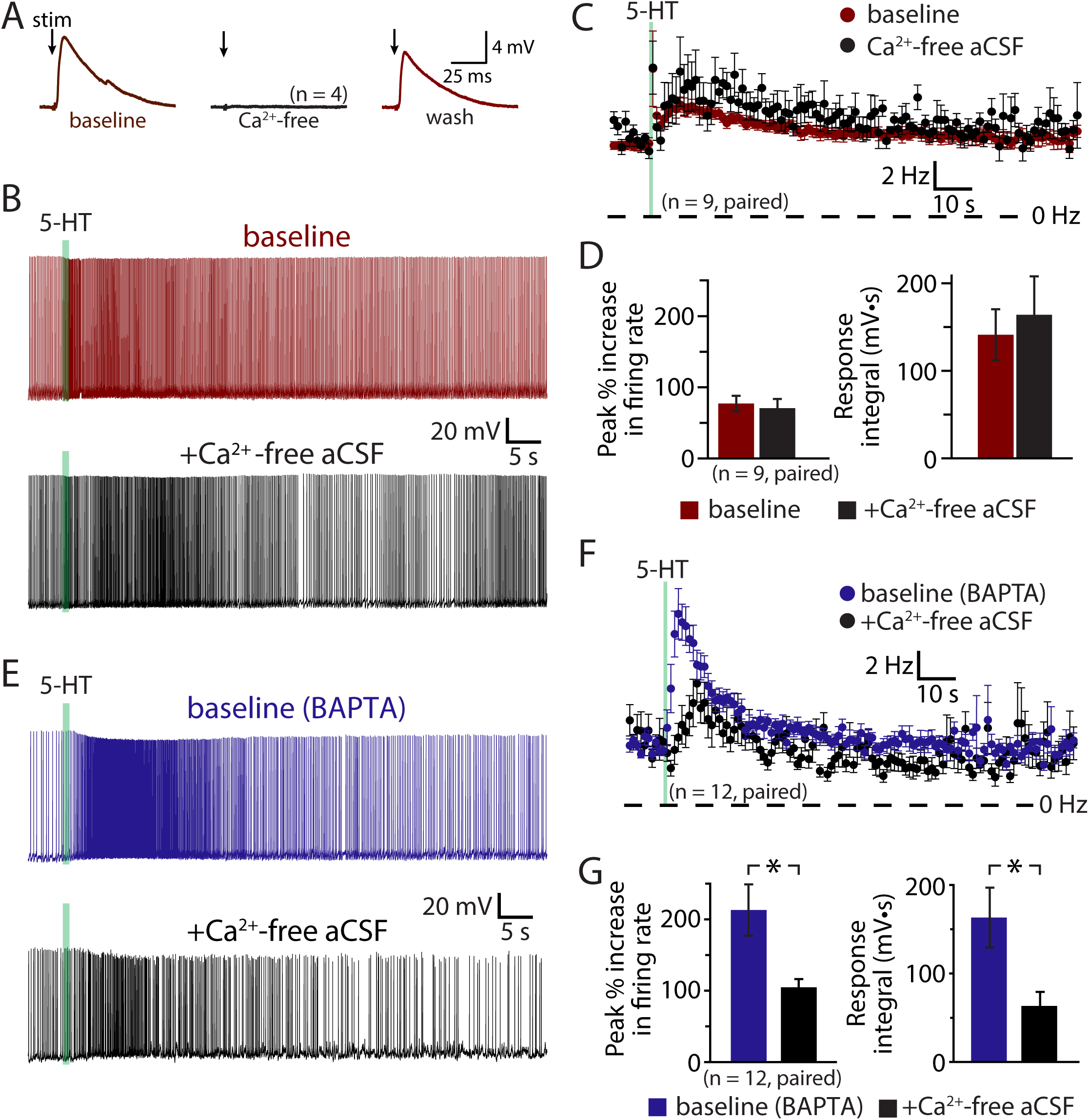
Serotonergic excitation involves a calcium conductance. **(A)** Electrically evoked EPSPs (indicated by arrows) in a COM neuron in baseline conditions (red), after replacement of extracellular calcium with magnesium (black), and after return to normal calcium conditions (red). **(B)** Serotonergic responses in a COM neuron in control conditions (red trace) and after replacement of extracellular calcium with magnesium (black trace). Green bars represent time of 5-HT application. **(C)** Plots of mean instantaneous spike frequencies in baseline (red) and zero-calcium (black) conditions for 9 COM neurons. **(D)** Comparisons of peak increases in instantaneous spike frequencies (left) and response integrals (right) for COM neurons before (red) and after (black) removal of extracellular calcium. **(E)** Responses to 5-HT (green bars) in COM neurons recorded with 10 mM BAPTA in the whole-cell pipette, before (blue) and after (black) removal of extracellular calcium. **(F)** Plots of mean (± SEM) instantaneous spike frequencies for 12 BAPTA-filled COM neuron in baseline (blue) and calcium-free (black) conditions. **(G)** Comparisons of 5-HT response amplitudes (left) and integrals (right) in BAPTA-filled COM neurons in baseline (blue) and calcium-free (black) conditions.

The lack of effect of extracellular calcium removal on 5-HT responses likely reflects competing cellular processes. For instance, even as removal of calcium may enhance some 5-HT-gated conductances, it will also eliminate the calcium-dependent ADP conductance and the calcium component of any calcium-permeable cation conductances contributing to excitation. To further explore the impact of extracellular calcium on serotonergic signaling in COM neurons, we filled additional neurons with BAPTA (10 mM) and measured serotonergic responses before and after replacement of extracellular calcium with magnesium (**Figure 4E**,**F**). As in our initial experiments (above), the presence of intracellular BAPTA led to robust serotonergic excitation in baseline (i.e., normal extracellular calcium) conditions, with peak increases in ISF being 213 ± 36% above initial firing rates, and with response integrals being 163 ± 34 Hz•s (n = 12). Subsequent removal of extracellular calcium greatly diminished both peak excitation (to 105 ± 11%; p = 0.007, paired Student’s t-test) and response integrals (to 63 ± 16 Hz•s; p = 0.004, **Figure 4G**). Because intracellular calcium signaling was blocked throughout these experiments with BAPTA, the reduction in response amplitudes and integrals observed after removal of extracellular calcium cannot be explained by effects on the ADP or other calcium-dependent conductances. Instead, these results point to involvement of a calcium-permeable cation conductance that under normal conditions is suppressed by intracellular calcium, but which is enhanced after chelation of intracellular calcium. Because responses in the presence of both intracellular BAPTA (which itself doubles 5-HT response amplitudes) and calcium-free aCSF (which in the absence of BAPTA has no effect on 5-HT responses amplitudes) results in responses comparable in size to those in control conditions (i.e., with normal intracellular and extracellular calcium levels), this calcium-sensitive calcium conductance appears to be balanced with the ADP and other calcium-dependent conductances.

### Role of M-current in serotonergic excitation

A classic ionic effector associated with G_q_-coupled receptors that may also exhibit calcium-sensitivity is the M-(“muscarinic”) current mediated by voltage-dependent K_V_7 (KCNQ) potassium channels (Delmas and Brown, 2005). Does M-current contribute to serotonergic excitation of COM neurons? If so, does intracellular calcium suppress the contribution of M-current to excitatory responses? To test the role of M-current suppression in generating serotonergic excitation, we measured 5-HT responses in COM neurons before and after bath applying the selective M-current blocker XE991 (10 *μ*M; **Figure 5A**,**B**). XE991 depolarized RMPs, from -80 ± 2 to -76 ± 1 mV (n = 10; p = 0.014, paired Student’s t-test), and increased input resistances (R_N_), from 155 ± 14 to 185 ± 16 MΩ (p = 0.0042, paired Student’s t-test), indicating that the M-current contributes to the resting membrane conductance of layer 5 COM neurons in the mouse mPFC. XE991 also reduced both the magnitude (by 49 ± 5%) and integral (by 56 ± 15%) of serotonergic excitation. Peak increases in ISF in response to 5-HT dropped from 94 ± 15% in baseline conditions to 50 ± 11% after application of XE991 (p < 0.001, paired Student’s t-test), while response integrals dropped from 132 ± 23 Hz•s to 48 ± 16 Hz•s (p = 0.009, **Figure 5E**,**F**). The substantial reduction in response amplitudes and integrals in the presence of XE991 suggests that suppression of the M-current is a key mediator of serotonergic excitation in COM neurons.

**Figure 5.**
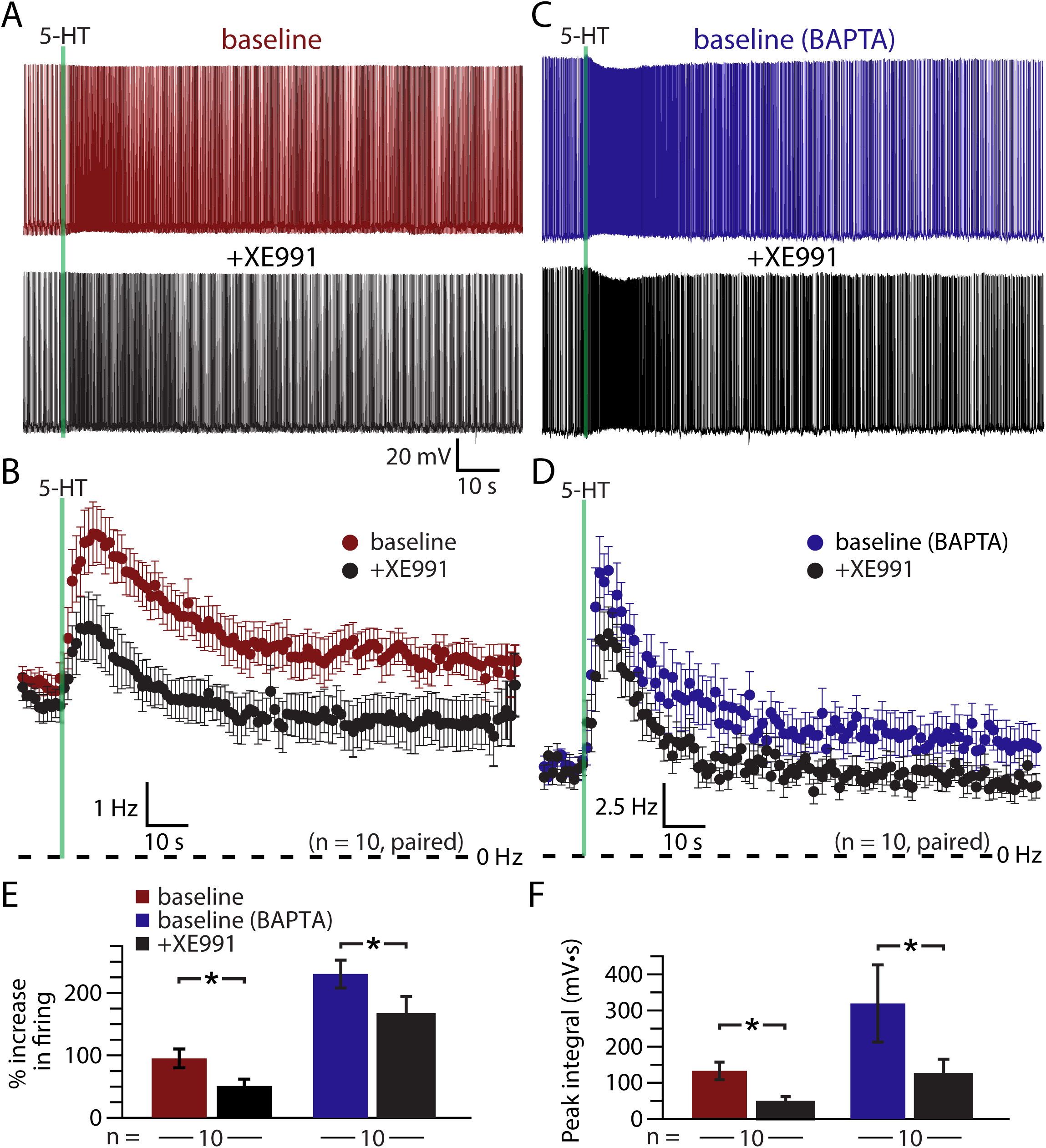
Suppression of K_V_7 channels contributes to serotonergic excitation of COM neurons. **(A**,**C)** Responses to 5-HT (green bars) in a control COM neuron (red, **A**) and a COM neuron patched with 10 mM BAPTA (blue, **C**) in baseline conditions and after blockade of K_V_7 channels with 10 μM XE991 (black). **(B**,**D)** Plots of mean instantaneous spike frequencies for control (**B**) and BAPTA-filled (**D**) COM neurons in baseline (red or blue) and XE991 (black) conditions. **(E**,**F)** Comparisons of peak increases in instantaneous spike frequencies (**E**) and response integrals (**F**) in baseline conditions (red or blue) and after addition of 10 μM XE991 (10 μM; black) for control and BAPTA-filled COM neurons.

If calcium sensitivity of M-current (see, for instance, Selyanko and Brown, 1996) contributes to the enhanced serotonergic excitation observed after intracellular calcium chelation, 5-HT responses after blockade of K_V_7 channels should be similar in both control and BAPTA-filled neurons. We tested this prediction by recording from a second group of neurons (n = 10) filled with 10 mM BAPTA (**Figure 5C**,**D**). Consistent with our earlier experiments using intracellular BAPTA, baseline responses to 5-HT were much larger in amplitude (mean increase in ISF was 230 ± 22%; p < 0.001 vs neurons lacking BAPTA; Student’s t-test, n = 10 per group), and integral (319 ± 107 Hz•s; p = 0.10) in BAPTA-filled neurons relative to neurons with intracellular calcium signaling intact. However, intracellular BAPTA did not enhance the impact of K_V_7 blockade with 10 μM XE991, in which response amplitudes (+164 ± 30% above baseline values) remained three times larger than in control neurons treated with XE991 (p = 0.002, Student’s t-test; **Figure 5E**,**F**). However, mean integrals (122 ± 42 Hz•s), while generally larger than in control neurons treated with XE991, were not statistically different in magnitude (p = 0.12). Overall, peak response amplitudes were reduced by 29 ± 9% (p = 0.022 vs baseline responses; paired Student’s t-test) and integrals by 57 ± 11% (p = 0.024 vs baseline responses) in BAPTA-filled neurons. Thus, the XE991-sensitive portions of serotonin responses in control and BAPTA recording conditions were similar in control and BAPTA conditions, suggesting that M-current alone does not account for the calcium-sensitivity of 5-HT excitation in COM neurons.

The results above suggest that 5-HT acts via at least three distinct mechanisms (K_V_7 suppression, the ADP conductance, and a calcium-sensitive calcium conductance) to enhance the excitability of COM neurons. To test whether M-current is the dominant potassium conductance contributing to serotonergic excitation, we enhanced the driving force for potassium six-fold by lowering the external potassium concentration ([K^+^]_o_) to 0.5 mM (replaced with equimolar sodium; **Figure 6**). This manipulation, which increases the driving force for potassium, will enhance the impact of M-current suppression by 5-HT, but will also act to reduce the net current through potassium-permeable nonspecific cation conductances. In neurons patched with control intracellular solution, lowering [K^+^]_o_ revealed a brief inhibition occurring immediately after 5-HT application, which was absent in control conditions (**Figure 6A**,**C**). While not pharmacologically challenged, these inhibitory responses appeared identical to G_q_-driven hyperpolarizations mediated by SK-type potassium channels that commonly occur in pyramidal neurons in response to M1 muscarinic receptor activation (Gulledge et al., 2009), but which are only rarely observed in response to 5-HT in control conditions. Lowering [K^+^]_o_ enhanced this early potassium conductance, and reduced the magnitude of serotonergic excitation by 31 ± 9% (n = 10, paired). In control conditions (e.g., 3 mM [K^+^]_o_), 5-HT generated peak responses of 82 ± 15% with integrals of 157 ± 44 Hz•s. After reducing extracellular potassium to 0.5 mM, peak excitation was 61 ± 15% (p = 0.003 relative to control conditions) with integrals of 117 ± 47 Hz•s (p = 0.0573, **Figure 6D**). Because the enhanced driving force for potassium is expected to increase the contribution of M-current suppression, the small reductions in response magnitudes and integrals suggest participation of potassium-permeable nonspecific cation conductances, such as the ADP conductance (Haj-Dahmane and Andrade, 1998).

**Figure 6.**
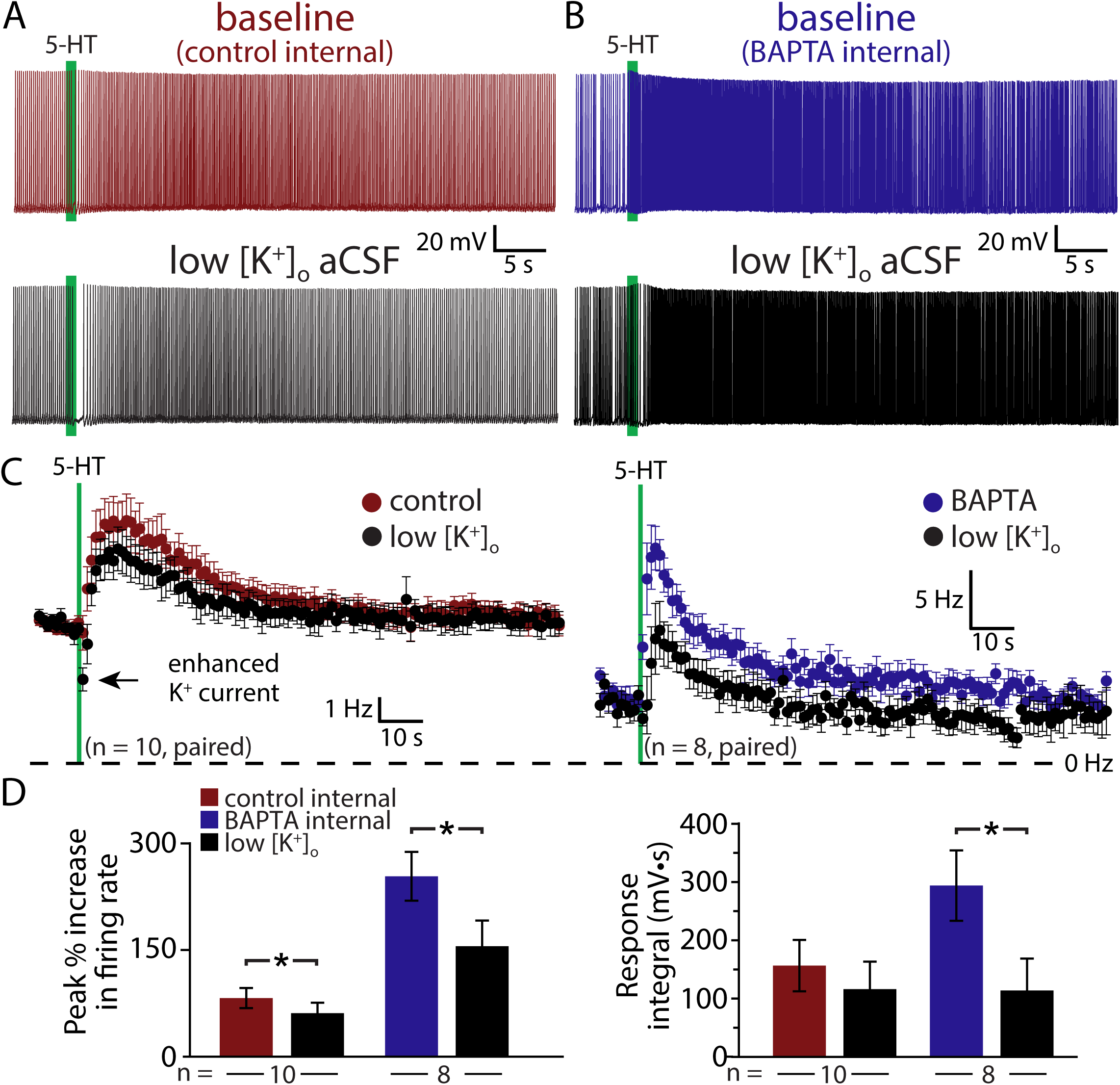
A role for non-selective cation conductances. **(A**,**B)** Responses to 5-HT (green bars) in baseline conditions and after reducing extracellular potassium to 0.5 mM (black traces) in control (red, **A**) or BAPTA-filled (10 mM, blue, **B**) COM neurons. **(C)** Plots of mean instantaneous spike frequencies for COM neurons in baseline (red or blue) and low [K^+^]_o_ (black) conditions for neurons patched with control intracellular solution (left) or with BAPTA included in the pipette solution (right). **(D)** Comparisons of peak increases in instantaneous spike frequencies (left) and response integrals (right) in baseline conditions (red or blue) and after lowering [K^+^]_o_ to 0.5 mM (black).

To further test the impact of potassium manipulations on 5-HT responses in the absence of the ADP nonspecific cation conductance and calcium-activated potassium channels, we repeated experiments in COM neurons filled with 10 mM BAPTA (**Figure 6B**,**C**). With intracellular calcium chelated, baseline responses were again large, with 254 ± 34% increases in ISF and 295 ± 61 Hz•s response integrals (n = 8). When [K^+^]_o_ was lowered to 0.5 mM, response amplitudes were reduced by 42 ± 7% (to 156 ± 36% above baseline firing rates; p < 0.001; paired Student’s t-test), while integrals were decreased by 63 ± 12% (to 114 ± 55 Hz•s; p = 0.029; **Figure 6D**). The percent reductions in both amplitudes (p = 0.342) and integrals (p = 0.559) after lowering [K^+^]_o_ in BAPTA filled neurons were similar to those in control neurons, suggesting that the calcium-sensitive conductance enhanced by intracellular BAPTA must also be a potassium-permeable nonspecific cation conductance.

## Discussion

Activation of G_q_-coupled receptors, including serotonergic 2A and muscarinic M1 receptors, enhances the intrinsic excitability of many neurons throughout the nervous system. As ubiquitous and robust as these responses are, it is surprising that there is not yet consensus on their underlying mechanisms. Data regarding cortical 2A signaling are particularly limited, in part due to the selective expression of these receptors in subpopulations of cortical neurons in adult animals (Weber and Andrade, 2010; Avesar and Gulledge, 2012). Therefore, much of our understanding of 2A receptor signaling has been inferred from studies of the more ubiquitous G_q_-coupled muscarinic ACh receptors. Here, we selectively targeted 2A-responsive COM neurons to probe the mechanisms underlying serotonergic excitation. Our results confirm that 2A receptors and muscarinic receptors share downstream signaling pathways, and point to at least three distinct ionic effectors contributing to serotonergic excitation: suppression of K_V_7 conductances, activation of a nonspecific cation channel that is both calcium-sensitive and calcium-permeable, and activation of the calcium-dependent nonspecific cation conductance responsible for the ADP (**Figure 7**).

**Figure 7.**
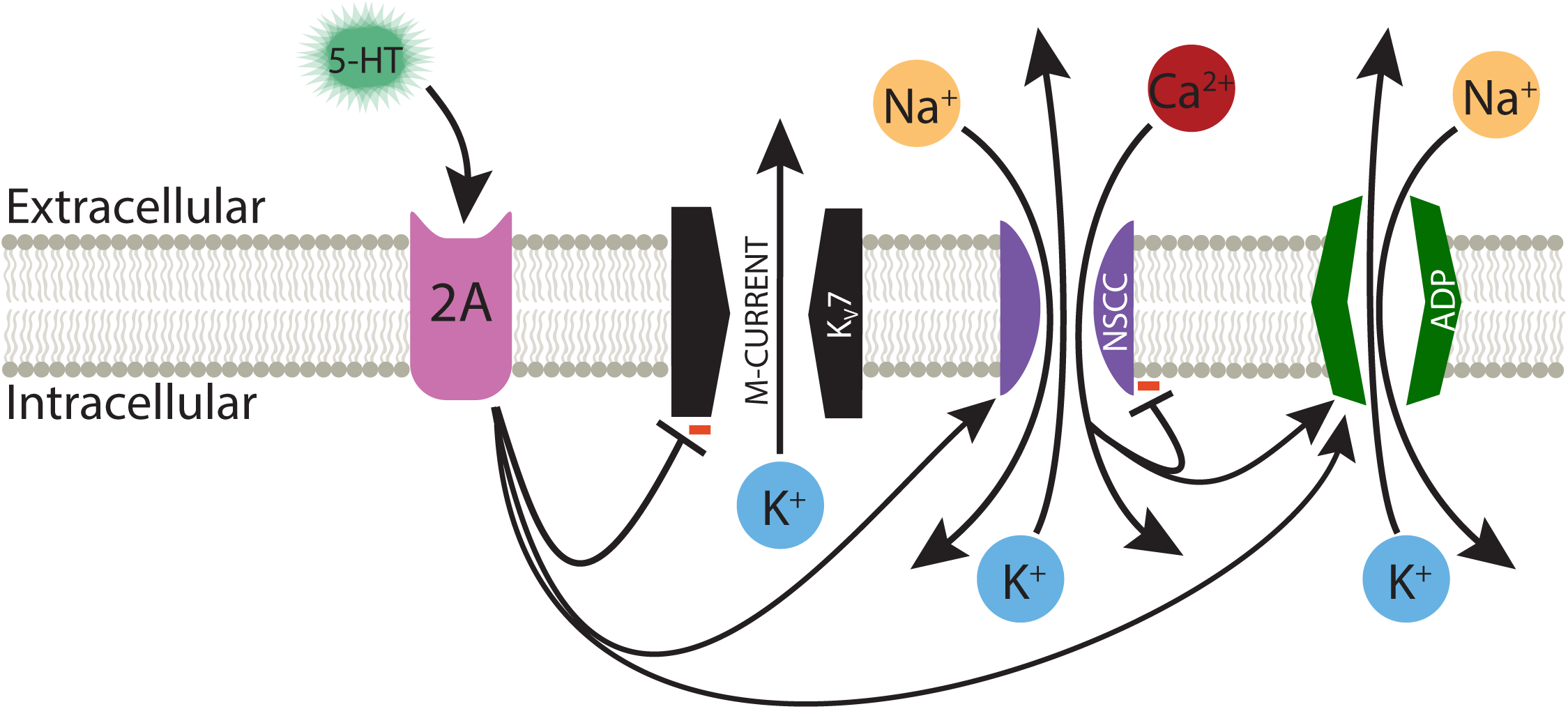
Ionic effectors contributing to serotonergic excitation of COM neurons. Activation of G_q_-coupled 5-HT_2A_ receptors in COM neurons targets a range of ionic effectors, including suppression of M-current, calcium-dependent activation of the ADP current, and activation of a calcium-sensitive, but calcium-permeable, nonspecific cation conductance (NSCC).

### Ionic conductances contributing to serotonergic excitation of COM neurons

Both M-current suppression (Caulfield et al., 1994) and activation of nonspecific cation conductances (Haj-Dahmane and Andrade, 1996) have been proposed as mechanisms for G_q_-mediated excitation. We found that under control conditions, with intracellular calcium signaling intact, suppression of M-current accounted for about half of the total serotonergic response. While K_V_7 channels are reported to be calcium-sensitive (Selyanko and Brown, 1996), the magnitude of M-current suppression was similar in control neurons and those filled with BAPTA, with 5-HT responses in the presence of both BAPTA and XE remaining much larger than responses measured in control conditions with intracellular calcium signaling and M-current intact. This suggests that the calcium-sensitivity of K_V_7 channels cannot not account for the larger 5-HT responses observed after chelation of intracellular calcium.

We also confirmed that 5-HT generates calcium-dependent ADPs in COM neurons. However, quantifying the relative contribution of the ADP conductance to enhanced action potential generation is problematic, as the ion channel mediating the ADP remains a matter of debate (Kuzmiski and MacVicar, 2001; Yan et al., 2009; Dasari et al., 2013; Lei et al., 2014), and attempts to block the conductance with intracellular BAPTA, or by removal of extracellular calcium, paradoxically enhanced the magnitude of serotonergic excitation. Indeed, in four independent experimental groups, baseline responses to 5-HT in the presence of intracellular BAPTA were more than twice as large as responses in control neurons with normal intracellular calcium signaling. This finding demonstrates that the ADP conductance is not required for serotonergic excitation, and that the net effect of normal levels of intracellular calcium is moderation, rather than amplification, of serotonergic excitation.

Two observations suggest that the calcium-sensitivity of 5-HT responses reflects a nonspecific cation conductance that itself is calcium permeable. First, the involvement of nonspecific cation conductances is strongly suggested by the effect of reducing extracellular potassium on 5-HT excitation. While this manipulation enhances selective potassium currents (e.g., the M-current), it also lowers the equilibrium potential for potassium-permeable nonspecific cation conductances. Thus, the reduction in serotonergic excitation observed after lowering extracellular potassium suggests the involvement of nonspecific cation conductances, such as the ADP conductance (Haj-Dahmane and Andrade, 1998). Since we continued to observe a reduction in response amplitudes after ADP currents were blocked by BAPTA, the remaining calcium-sensitive calcium conductance, which was amplified by chelation of intracellular calcium, is likely also a potassium-permeable nonspecific cation conductance. Second, we found that the effect of removing extracellular calcium was dependent on the baseline state of intracellular calcium signaling within neurons. In control conditions, with calcium-signaling intact, removal of extracellular calcium had no impact on 5-HT response amplitudes or integrals. Yet, when intracellular calcium signaling was blocked with BAPTA, and baseline responses to 5-HT were twice as large as in control neurons, removal of extracellular calcium greatly reduced the amplitudes and integrals of 5-HT responses such that they became comparable in size to control 5-HT responses observed in the absence of BAPTA. These findings suggest the involvement of a calcium-permeable nonspecific cation conductance that is itself suppressed by intracellular calcium (**Figure 7**). Under normal conditions, with intracellular calcium signaling intact, removal of extracellular calcium should have two opposing effects: it would eliminate a charge carrier for the calcium-permeable nonspecific cation conductance, thereby hyperpolarizing the effective equilibrium potential for the conductance, but would also enhance the maximum conductance by relieving the channel from negative regulation by intracellular calcium. Our results suggest these two competing influences are fairly balanced with the calcium-dependent ADP conductance, such that calcium-removal in control conditions has little impact on serotonergic responses. However, when the conductance is maximized by chelating intracellular calcium with BAPTA, subsequent removal of extracellular calcium acts only to reduce net current, such that 5-HT responses become smaller and equivalent to responses observed in control conditions.

These findings regarding calcium regulation of serotonergic excitation are most comparable to those of Magistretti et al. (2004), who found G_q_-mediated cholinergic responses in layer 2 pyramidal neurons in the entorhinal cortex to be bidirectionally influenced by calcium influx. They found that modest increases in intracellular calcium enhanced net inward current (attributed to a calcium-dependent ADP-like conductance), while larger increases in intracellular calcium, including normal levels of calcium accumulation during periods of action potential generation, acted to suppress cholinergic excitation (Magistretti et al., 2004). While Magistretti et al. concluded that their results reflected bidirectional calcium regulation of a the ADP conductance, results from our experiments using BAPTA and calcium-free aCSF dissociate the calcium-activated ADP conductance from the additional calcium-sensitive nonspecific cation conductance. Co-activation of two nonspecific cation conductances with opposing calcium regulation may contribute to the robustness of G_q_-mediated excitation by allowing it to maintain a consistent net current amplitude independent of intracellular calcium levels, while also promoting calcium-influx when intracellular calcium levels are low. This negative feedback of intracellular calcium on the calcium-permeable conductance should act to stabilize intracellular calcium levels, and could provide a mechanism for calcium store refilling after G_q_ receptor signaling (Dasari et al., 2017).

### G_q_-signaling in cortical projection neurons

The ionic mechanisms contributing to serotonergic excitation of COM neurons appear to be conserved across cortical neuron subtypes and transmitter signaling systems. In a recent preprint (Baker et al., 2017), we report that G_q_-mediated cholinergic excitation occurs preferentially in corticopontine (CPn) neurons in the mouse mPFC (see also Dembrow et al., 2010), and involves the same combination of ionic effectors (M-current suppression, the ADP conductance, and a calcium-sensitive and calcium-permeable non-specific cation conductance) as described here in relation to serotonergic 2A signaling in COM neurons. However, some notable differences exist in G_q_-mediated excitation of COM and CPn neurons by 5-HT and ACh, respectively. For instance, although stimulation of muscarinic ACh receptors robustly generated transient SK-channel mediated inhibition in both COM and CPn neurons (Baker et al., 2017), we rarely observed SK-dependent inhibition in mouse COM neurons following 5-HT application, even as ACh and 5-HT generate similar increases in ISF (∼100% above baseline levels) in these same neurons. Even after enhancing the driving force for potassium channels with lowered extracellular potassium, SK-like responses to 5-HT were observed in only 7 of 10 neurons, and had shorter durations (0.7 ± 0.2 s) than SK responses generated by ACh in either COM (∼1.5 s) or CPn (∼ 1 s) neurons (Baker et al., 2017). This may reflect differential subcellular localization of M1 muscarinic receptors relative to serotonergic 2A receptors in COM neurons, or subtle differences in G_q_-coupling to second messenger systems. Another difference is the relative contribution of ionic effectors in generating excitatory responses. In CPn neurons, suppression of M-current accounted for only ∼20% of total cholinergic excitation, while chelation of intracellular calcium enhanced responses by only ∼25% (Baker et al., 2017). This suggests that the ADP conductance plays an overall larger role in mediating cholinergic excitation in CPn neurons relative to COM neurons. Thus, M-current suppression and the calcium-sensitive nonspecific cation conductance appear more important for serotonergic excitation of COM neurons, relative to cholinergic excitation of CPn neurons. Since CPn neurons in mice do not exhibit serotonergic excitation, future studies that directly compare the mechanisms of cholinergic and serotonergic excitation in COM neurons will be necessary to determine whether these signaling differences are cell-type-or transmitter-dependent.

## Author contributions

AG and ES designed research. AG, AB, and ES conducted research and analyzed data. AG, ES, and AB wrote the manuscript.

## Acknowledgements

The authors thank Saiko Ikeda for technical assistance.

